# Effect of Zn fertilization on yield, protein content, protein yield and zinc use efficiency in pearl millet

**DOI:** 10.1101/2022.12.08.519627

**Authors:** Ganesh Narayan Yadav, Rakesh Summaria, Shankar Lal Yadav, Seema Sharma, L. R. Yadav

## Abstract

Two year field experiment was conducted at Rajasthan Agriculture Research Institute, Durgapura, during rainy seasons of 2019 and 2020 in factorial block design with three replications to study the growth, productivity and economics of pearl millet – chickpea cropping system by using three genotypes (RHB-173, MPMH-17 and RHB-177) and five treatments of zinc [0, 5, 7.5 kg Zn/ha through ZnSO_4_.7H_2_O and 5, 7.5 kg Zn/ha through Zn-EDTA) applied to pearl millet and two foliar spray treatments applied in both of crop at booting stage in pearl millet and pre-flowering stage in chickpea. Results revealed that total nitrogen and zinc contents of chickpea were increased significantly and highest with the residual effect of 7.5 kg Zn/ha through Zn-EDTA and 7.5 kg Zn/ha through ZnSO_4_.7H_2_O and phosphorus contents of chickpea was recorded depressive effect on residual zinc fertilization, while, potassium contents was not influenced significantly. However, the significant increase in uptake of nutrients nitrogen, phosphorous, potassium and zinc were obtained with 7.5 kg Zn/ha through Zn-EDTA higher per cent over control, during the years 2019-20 and 2020-21. The highest nitrogen, protein content and zinc contents and uptake values were obtained with 5, 7.5 kg Zn/ha through Zn-EDTA. The residual effect of 7.5 kg Zn/ha through Zn-EDTA was at par with 7.5 kg Zn/ha through ZnSO_4_.7H_2_O. The increased in total uptake of nitrogen, it could be ultimately increased protein yield were 16.54 and 27.48 per cent, during respective years over control. The highest nutrient use efficiency was obtain with lowest of fertilizer dose of Zn-EDTA as compare to zinc sulphate.

## Introduction

In India, pearl millet is one of the important millet crops which flourishes well even under adverse conditions. It is nutritious and palatable and adapted to drought and poor soil fertility, but responds well to good management and higher fertility levels.

Since zinc is growth promoting substance which plays indispensable role in various plant physiological processes (Khinchi *et al*., 2017). A greater part of soil phosphorus and zinc are in the form of insoluble phosphates that can be utlized by using zinc EDTA, its higher solubility efficiency as compare to zinc sulphate, the remaining fixed unavailable zinc will be available and that residue fertilizer can be utilize by succeeding crop. Barman *et al*. (1998) were of the opinion that residual effect of different levels of zinc applied to soybean crop significantly increased dry matter of succeeding wheat crop. The Zn-EDTA was observed higher zinc use efficiency as compared to zinc sulphate.

## Materials and Methods

A field experiment was conducted, during rainy seasons of the years 2019 and 2020 at research farm of Rajasthan Agricultural Research Institute-Durgapura (SKNAU, Jobner). The soil of the field was loamy sand, low in available nitrogen and phosphorous, medium in potassium and low in zinc content (138.5, 25.1, 181 kg/ha and 0.36 ppm) during 2019 and (135.2, 23.2, 180 kg/ha and 0.34 ppm) during 2020, respectively in 0-30 cm soil depth. The soil pH was 8.4 and 8.2 and per cent organic carbon content was 0.16 and 0.13 per cent with respective years. Treatments comprised of thirty treatment combinations consisting of three genotypes (RHB-173, RHB-177 and MPMH17), five zinc treatments (0, 5, 7.5 kg Zn/ha through Zn-EDTA and 5, 7.5 kg Zn/ha through ZnSO_4_.7H_2_O) and two foliar spray (No spray and 0.5 % zinc sulphate foliar spray at booting stage) were tested in factorial block design with three replications. A uniform dose of 70 kg N/ha and 40 kg P/ha along with zinc as per treatment were drilled through zinc sulphate (21%) and Zn-EDTA (12%), respectively. To compensate the sulphur obtained from different levels of zinc compensatory dose of sulphur applied through elemental sulphur. Not a major insect/disease was observed during the life cycle of pearl millet in the experiment, but weeds were manually controlled once (30 days after sowing). Economics of treatments were worked out using market price of inputs and minimum support price of outputs.

The plant samples were analysed for estimation of N with methods of Snell and Snell, 1949; P and K with Jackson, 1973 and Zn contents through the method Lindsay and Norvell, 1978. The per cent crude protein in grain and stover was computed by multiplying the nitrogen content in grain and straw with a constant factor of 6.25 (A.O.A.C., 1960).

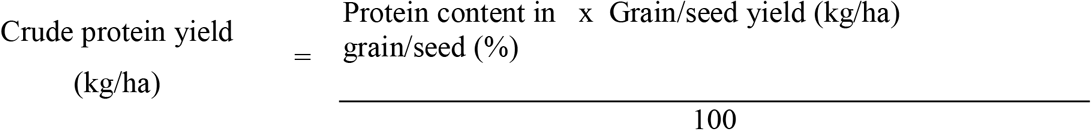

The nutrient use efficiency for zinc were worked with the following formula given by Craswell and Godwin (1984) and expressed in kg grain or seed / kg nutrient.

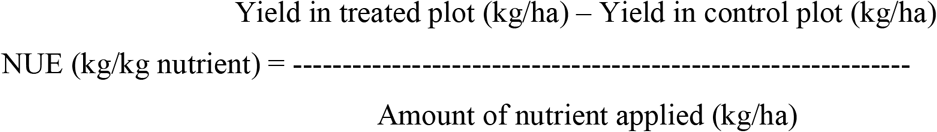

## Result and Discussion

### Crude protein content

Among the pearl millet genotypes there was found significant improvement with respect to protein contents with the genotype RHB-173 over RHB-177 and MPMH17.

The total protein contents of pearl millet were significantly increased with different levels of zinc over control during 2019 and 2020, respectively. The application of Zn-EDTA was found superior over ZnSO_4_.7H_2_O. The highest protein contents was found with the application of 7.5 kg Zn/ha through Zn-EDTA followed by with the application of 7.5 kg Zn/ha through ZnSO_4_.7H_2_O. The increase in yields of pearl millet owing to effects of zinc levels might be due to the fact that boost photosynthetic rate and enhanced absorption and nutrition of zinc so accumulation of proteins contents in crop.

Nourishment with zinc helps in improving nitrogen content of plant through biological nitrogen fixation (BNF) though, nitrogen appears to be synergistic with potassium and zinc, which may leads to increase in many physiological and molecular activities which in turn improve yield attributing characters (Cakmak *et al*., 2010). In dry land areas zinc application increases absorption of minerals by roots (Singh *et al*., 2017).

### Crude protein yield

During 2019 and 2020 increasing level of zinc significantly enhanced the yield then ultimately increased of protein yield by crop as compared to the control and the highest values of crude protein yield were recorded with the application of 7.5 kg Zn/ha through Zn-EDTA over control, however, it was at par with all other zinc treatments. The highest crude protein yield was recorded with the application of 7.5 kg Zn/ha through Zn-EDTA, followed by 7.5 kg Zn/ha through ZnSO_4_.7H_2_O, 5 kg Zn/ha through Zn-EDTA, 5.0 kg Zn/ha through ZnSO_4_.7H_2_O during both year. Biological nitrogen fixation increased by zinc, thus the availability of nitrogen in the rhizosphere is increased and that in turn improved its uptake. (Singh et al. 2015) observed that, Zn increased the cation exchange capacity of roots which might have helped in increased absorption of zinc from the soil and consequently increased protein yield in pearl millet. The increase in zinc could be attributed to synergistic effect between nitrogen and zinc and owing to the positive interaction of crude protein yield respectively. The present findings support the results of Ashoka *et al*. (2008), Morshedi and Farahbakhsh (2010).

### Zinc use efficiency

Effect of 5 kg Zn/ha through Zn-EDTA obtained highest zinc use efficiency of pearl millet and it was superior over control, however it was at similar effect with the 7.5 kg Zn/ha through ZnSO_4_.7H_2_O, 5 kg Zn/ha through Zn-EDTA and 5 kg Zn/ha through ZnSO_4_.7H_2_O, during both the years. During 2019 and 2020, effect of 0.5 % zinc sulphate foliar spray at booting was found observed the higher zinc use efficiency, during both the years. Positive improvement in Zn-EDTA due to fertilization with zinc that was augmented by coupling with zinc had a contributory effect in improving yield attributes. Photosynthetic reactions depend on an adequate supply of zinc because of the presence of this metal in key photosynthetic enzymes, such as RuBisCO (ribulose-1,5-bisphosphate carboxylase / oxygenase) and carbonic anhydrase (in C_4_ plants) Kryvoruchko (2017).

## Conclusion

On the basis of this study it is concluded that the effective improvement was observed in increasing crude protein content, crude protein yield and zinc use efficiency of chickpea crop through residual zinc fertilization of 5 kg Zn/ha through Zn-EDTA and 5 kg Zn/ha through ZnSO_4_.7H_2_O.

---

**Table 1.**
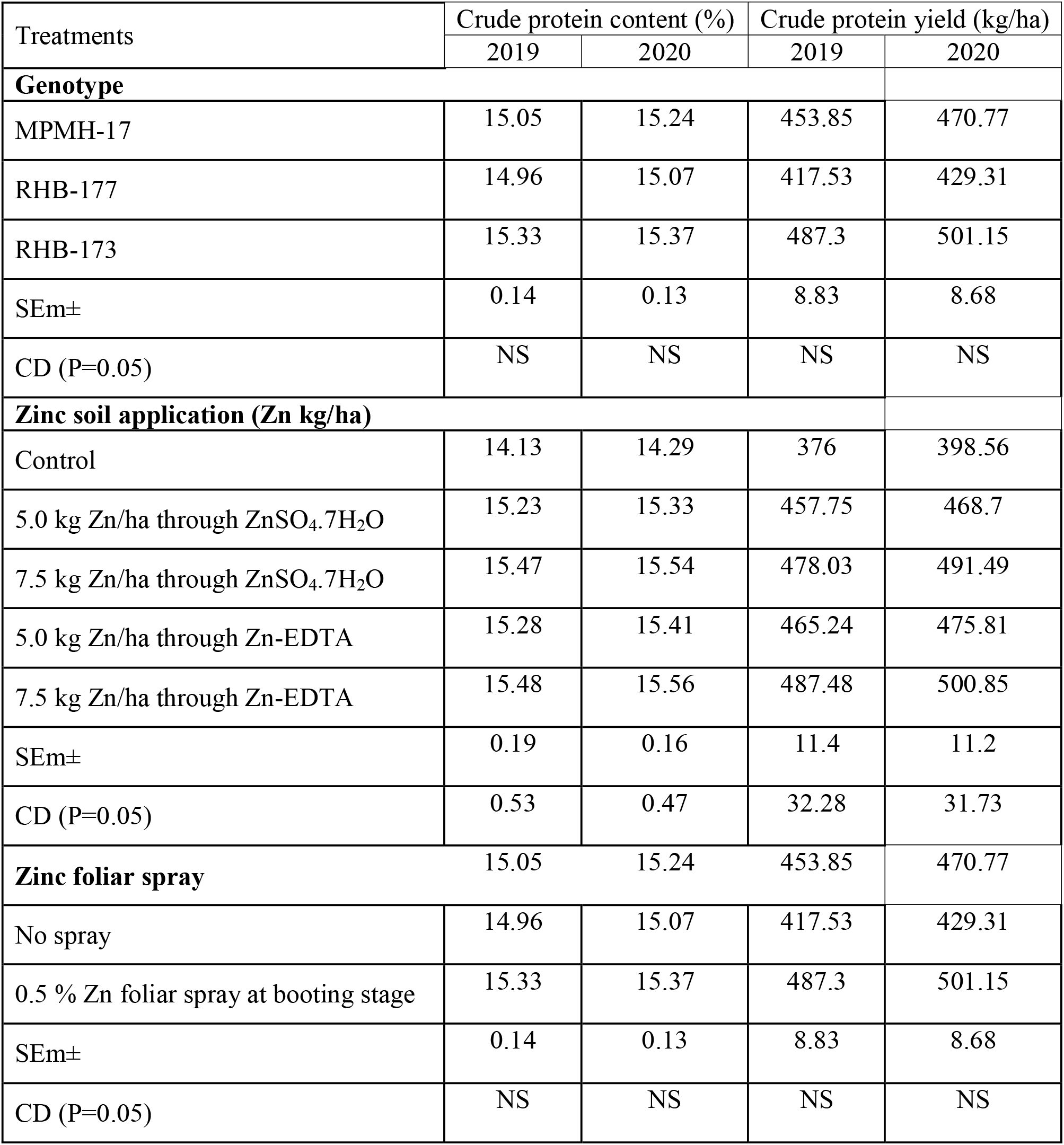
Effect of zinc fertilization on crude protein contents and crude protein yield of pearl millet during both the years.

**Table 2.** Effect of zinc fertilization on zinc use efficiency of pearl millet during both the years.

| Treatment | ZUE(kg/kg) |  |  |
| --- | --- | --- | --- |
|  | 2019 | 2020 | Pooled |
| **Zinc soil application (Zn kg/ha)** |  |  |  |
| Control | 0.00 | 0.00 | 0.00 |
| 5.0 kg Zn/ha through ZnSO<sub>4</sub>.7H<sub>2</sub>O | 140.79 | 125.32 | 133.05 |
| 7.5 kg Zn/ha through ZnSO<sub>4</sub>.7H<sub>2</sub>O | 129.45 | 125.31 | 127.38 |
| 5.0 kg Zn/ha through Zn-EDTA | 164.54 | 146.52 | 155.53 |
| 7.5 kg Zn/ha through Zn-EDTA | 140.36 | 139.63 | 139.99 |
| SEm± | 30.88 | 27.53 | 20.69 |
| CD (P=0.05) | - | - | - |
| **Zinc foliar spray** |  |  |  |
| No spray | 108.24 | 78.20 | 93.22 |
| 0.5 % Zn foliar spray at booting stage | 121.82 | 136.50 | 129.16 |
| SEm± | 19.53 | 17.41 | 13.08 |
| CD (P=0.05) | - | - | - |

**Fig. :1.**
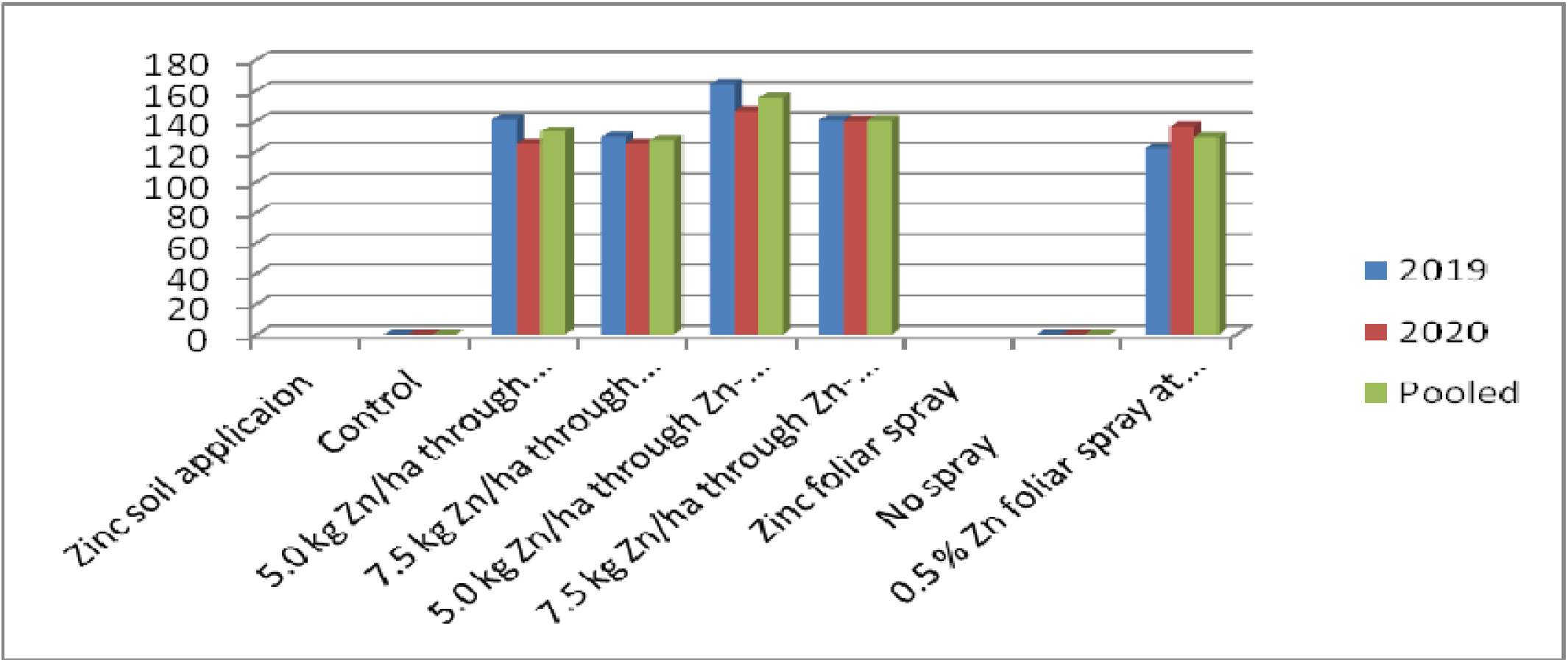
Zinc use efficiency influenced by Zn fertilization.

## References

Barman K K, Ganeshamurthy A N, Takkar P N. Zinc requirement of soybean-wheat cropping sequence in some swell-shrink soils. Indian journal of Agricultural science 1998; 68 (12): 759–61.

Cakmak I, Pfeiffer W H, Mc Clafferty B. Bio fortification of durum wheat with zinc and iron. Cereal Chem 2010; 87: 10–20.

Jackson M L. Soil Chemical Analysis. Prentice Hall of India Pvt. Ltd., New Delhi 1973.

Khinchi V, Kumawat S M, Dotaniya C K. Effect of Nitrogen and Zinc Levels on Yield and Economics of Fodder Pearl Millet (Pennisetum americanum L.). International Journal of Pure and Applied Bioscience. 2017; 5(3):426–430.

Kryvoruchko I S. Zn-use efficiency for optimization of symbiotic nitrogen fixation in chickpea (Cicer arietinum L.). Turkish Journal of Botany 2017; 41: 423–441

Kumawat S, Sammauria R, Kumawat P. Growth and productivity of succeeding pearl millet (Pennisetum glaucum) influenced by residual phosphorus, zinc and zinc solubilizer International Journal of Chemical Studies 2018; 6(5): 272–275

Lindsay WL, Norwell WA. Development of DTPA soil test for zinc, iron, manganese and copper. Soil Science Society of America Journal 1978; 42: 421–428.

Singh L, Sharma P K, Jajoria M, Deewan P, Vermam R. Effect of Phosphorus and Zinc Application on Growth and Yield Attributes of Pearl millet (Pennisetum glaucum L.) under Rainfed Condition. Journal of Pharmacognosy and Phytochemistry 2017; 6 (1): 388-391

Snell FD, Snell CT. Colorimetric Methods of Analysis. 3rd Edn. Vol. II D. Van Nostrand Co. Inc., New York 1949.

